# Structural characterization of the PCV2d genotype at 3.3 Å resolution reveals differences to PCV2a and PCV2b genotypes, a tetranucleotide, and an N-terminus near the icosahedral 3-fold axes

**DOI:** 10.1101/614198

**Authors:** Reza Khayat, Ke Wen, Aleksandra Alimova, Boris Gavrilov, Al Katz, Jose M. Galarza, J. Paul Gottlieb

## Abstract

Porcine circovirus 2 (**PCV2**) is a T=1 non-enveloped icosahedral virus that has a major impact on the swine industry as an agent of porcine circovirus associate disease. PCV2 capsid protein sequences have been employed by others to provide a temporal description of the emerging genotypes. PCV2a is believed to be the earliest genotype and responsible for giving rise to PCV2b, which gives rise to PCV2d. The underlying mechanism responsible for the emerging genotypes is not understood. To determine if a change in the PCV2d capsid accompanies the emergence of this genotype, we determined the cryo-electron microscopy image reconstruction of PCV2d VLP at 3.3 Å resolution and compared it to the previously reported PCV2a and PCV2b structures. Differences between the CD and GH loops identify structural changes that accompany the emergence of PCV2b from PCV2a, and PCV2d from PCV2b. We also model additional amino acids for the N-terminus near the icosahedral 3-fold axes of symmetry and a tetranucleotide between the 5- and 2-fold axes of symmetry. To interpret the sequence diversity that defines the PCV2 genotypes on a structural platform we have performed structure-based sequence comparison. Our analysis demonstrates that each genotype possesses a unique set of amino acids located on the surface of the capsid that experience a high degree of substitution. These substitutions may be a response to the PCV2 vaccination program. The structural difference between PCV2a, b and d genotypes indicate that it is important to determine the PCV2 capsid structure as the virus evolves into different genotypes.

**Importance:** PCV2 is a significant epidemic agricultural pathogen that is the causative agent of a variety of swine illnesses. PCV2 infections have significant economic impact in the swine industry and must be controlled by vaccination. Outbreaks in farms vaccinated for PCV2 suggest that improvements to the current vaccination programs are needed. Better understanding of the assembly, structure, replication and evolution of these viruses is necessary for production of improved vaccines. The ability of PCV2 to rapidly shift genotypes suggests that expression systems capable of rapidly producing large quantities of virus-like particles should be pursued. To these ends we have established a mammalian cell-based virus-like particle expression system and performed high resolution structural studies of a new PCV2 genotype. Differences between the structure of this genotype and earlier genotypes demonstrate that it is important to study the PCV2 structure as it shifts genotypes.

## Introduction

Circoviruses are small nonenveloped icosahedral viruses that contain a circular, covalently closed, single stranded DNA (**ssDNA**) genome. Circoviruses have been identified in a variety of species and are known to cause infections in avian, aquatic, and terrestrial animals (1, 2). The variation of the capsid morphology and genome organization has recently led to the classification of the *Circoviridae* family into two separate genera, *Cyclovirus* and *Circovirus* (3). Genome sequences associated with the *Cyclovirus* genus has been identified from several vertebrate and invertebrate species, although the recognition of definitive host for this group is still unclear (2). The genus *circovirus* comprises porcine circoviruses types 1 (**PCV1**), 2 (**PCV2**), and 3 (**PCV3**) (2). PCV2 infections are responsible for significant mortality among swine as the causative agent of porcine circovirus associated disease (**PCVAD**) -weight loss, jaundice, stressed appearance, enlarged/depleted lymph nodes, pneumonia, enteritis, diarrhea, increased mortality and cull rates, abortions, stillbirths, and mummies (1, 4).

The PCV2 particle is approximately 19 nm in diameter, and the genome of PCV ranges from 1.7 kb to 2.3 kb in size. The circular nature of the genome has led to the viral family name *circovirus*. The evolutionary history has been explored and has allowed detailed phylogenetic trees and variations in the capsid surface structure to be described (5–8). The initial cryo-EM image reconstruction of several native circoviruses demonstrated that the capsid has *T=1* icosahedral symmetry (9). The PCV genome encodes for one structural capsid protein (**CP**). Expression and purification of the PCV2b CP protein from *E. coli* were demonstrated to self-assemble and mimic the overall morphology of the infectious virus. The crystal structure of the *E. coli* produced virus-like particle (**VLP**) visualized the CP fold to be that of the canonical viral jelly roll consisting of two four stranded β-sheets (10, 11). The loops connecting the β-strands form the features on the viral surface and may include the antigenic epitopes. The PCV2b CP was also expressed and purified as VLP from *Trichoplasia ni* insect cells (11). The cryo-EM image reconstruction of this VLP demonstrated that the N-terminus is located inside the capsid, and the authors concluded that the antigenic properties associated with the N-terminus is likely a result of the N-terminus transiently externalizing from the capsid via a process referred to as viral “breathing” (11–13). The externalization of the N-terminus may play an important role in the life cycle of the virus. While production of CP from *E. coli* and baculovirus expression systems have resulted in assembly of PCV2 VLPs that appear to resemble the structures of native PCV2 virus, the systems are quite different from a mammalian system which is a natural host for PCV2 virus. A mammalian expression system may provide better conditions for the examination of capsid assembly and structure; therefore, we have chosen HEK293 mammalian cells to evaluate the formation and structural characterization of the produced PCV2 VLPs.

The PCV2 CP entries in GenBank were recently categorized into eight different genotypes (PCV2a-h) (14). The chronological deposition of PCV2 sequences into GenBank suggests that PCV2a was the dominant global genotype until early-2000 when a genotype shift to PCV2b was observed (15, 16). The PCV2c genotype has only been reported from European countries and may have become extinct as there are only four depositions in GenBank. In 2013 Wei *et al*. reported and deposited a large quantity of PCV2d sequences into GenBank (5). Since then the number of PCV2d depositions in GenBank has significantly outpaced the deposition of PCV2b sequences (7, 14). The increase in the deposition of PCV2d sequences may be a result of PCV2d becoming an emerging and predominant genotype in Asia, Europe, North and South America. While the cause(s) for the increase in depositions remains unclear, the presence of PCV2d in vaccinated herds suggests that either the vaccine is not appropriately administered or that PCV2d represents a genotype resistant to vaccination (17, 18). Despite the significant number of phylogenetic studies of the PCV2 genotypes and the emerging importance of PCV2d on the global swine industry there are no reports describing the structure of the PCV2d capsid. To determine if a structural difference between PCV2b and PCV2d may explain the shift in genotypes, we established an expression system for producing large quantities of the PVC2d VLPs in mammalian cells (human embryonic kidney: HEK 293). Study of the cytoplasmic and nuclear fractions of the mammalian cells indicates that PCV2 assembles and remains in the nucleus within 72 hours of transfection. The cryo-EM image reconstruction of the VLPs determined to a resolution of 3.3 Å allows us to confidently model the atomic coordinates for amino acids 36-231. Comparison of the PCV2d atomic coordinates to that of PCV2a and PCV2b identifies loops CD and GH of PCV2b/d to significantly deviate from the PCV2a structure. These differences provide a structural explanation for the PCV2a to PCV2b shift in genotypes. Comparison of the coordinates for amino acids 36-45 to a recently reported PCV2 cryo-EM image reconstruction indicates a significant difference between the two models. We model the N-terminus to be near the icosahedral 3-fold axes of symmetry while Mo *et al*. model the N-terminus to be near the icosahedral 5-fold axes of symmetry (19). We discuss this discrepancy in the discussion section. We use the PCV2d cryo-EM image reconstruction to calculate ~1,200 nucleotides to be packaged into the VLP, and support this finding with absorption spectroscopy measurements. We further model a Pu-Pu-Py-Py tetranucleotide between the icosahedral 5-and 2-fold axes of symmetry. The strong density for the tetranucleotide suggests that the PCV2d capsid selects this nucleotide sequence from the pool of sequences present in the nucleus where capsid assembly occurs. We then compare the surface amino acids composition of PCV2a, b, d and determine that the amino acids that experience sequence diversity (identity) and variability (entropy) within each genotype are unique to that genotype. The PCV2a genotype experiences the greatest sequence diversity and the lowest sequence variability, whereas the PCV2b and d genotypes experience lower sequence diversity and greater variability. Expanding this analysis to include the remaining genotypes (1,278 unique entries) indicates that except for three amino acids (Met1, Pro15 and Arg147) every amino acid position has experienced a substitution. However, two groups of amino acids exhibit limited sequence variation and form distinct patches on the surface of the capsid. We propose that vaccines capable of directing antibodies to these patches may serve as universal vaccines for PCV2.

## Results

### Recombinant PCV2 VLPs assemble and remain in the nucleus of expression system

We sub-cloned the PCV2d capsid protein (**CP**) gene of C/2013/3 isolated in Taiwan (GenBank: AWD32058.1) into the mammalian expression plasmid pcDNA 3.4 in order to produce recombinant PCV2d VLP in mammalian cells (Fig. 1A). Transfection of suspension culture of HEK-293 mammalian cells with the plasmid results in the production of the CP and assembly of the VLP. To determine the location of VLP assembly (nucleus versus cytoplasm), we used immunofluorescence and Western blot analysis of transfected cells. The analysis was performed at three time points post-transfection: 24, 48, and 72 hours. The mouse anti-PCV2 capsid monoclonal antibody was used as a probe directed to amino acids 49-125 of the CP (exact epitope location is proprietary information of GeneTex). The transfected cells were subsequently lysed at each of the time points and three cellular fractions isolated, whole cell lysate (WCL), cytoplasm (CE) and nuclear (NE) extracts. At 48 hours immunofluorescence expressed CP becomes evident. At this time point CP was found in both the nucleus and the cytoplasm; by 72 hours the nuclear compartment appeared enriched (Fig 2). These results were confirmed by Western Blot analysis. These results demonstrate that while translation is, as expected, in the cell cytoplasm the protein is subsequently transported into the nucleus where capsid assembly occurs and the VLP remains. To date PCV2 VLP has been produced using insect cells, yeast, or *E. coli* (11), thus expression of VLP using mammalian cells provides an analogous substrate to that utilized by natural PCV2 infection.

**Figure 1.**
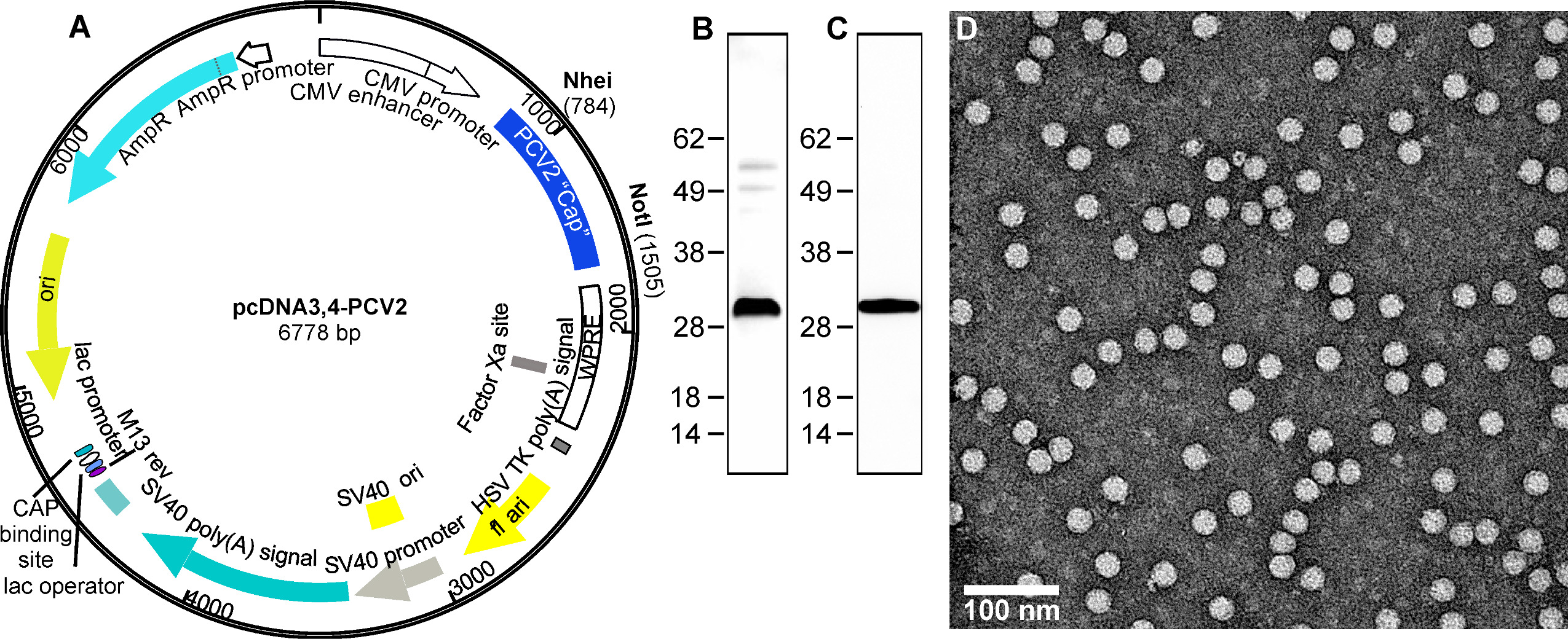
Expression of PCV2 virus-like particles in mammalian cells. A) Plasmid generated for expression of PCV2d capsid protein. The codon optimized PCV2d capsid gene was synthesized (Blue Heron Technologies, Bothell, WA) and cloned into expression vector pcDNA3.4 (Fisher Scientific). B) Protein expression was conducted in transiently transfected suspension cultures of Expi293 cells (Life Technologies). SDS-PAGE analysis of purified PCV2d VLPs (1 ug protein) and stained with Coomassie blue. C) SDS-PAGE analysis of purified PCV2d VLPs (0.5 ug protein) transferred to a nitrocellulose membrane and probed for a Western Blot with primary rabbit anti PCV2 capsid polyclonal antibody (Cab 183908, Abcam, UK). D) Negative stained electron microscopy micrograph of purified VLP stained with uranyl acetate. Particle sizes are approximately19 nm diameter.

**Figure 2.**
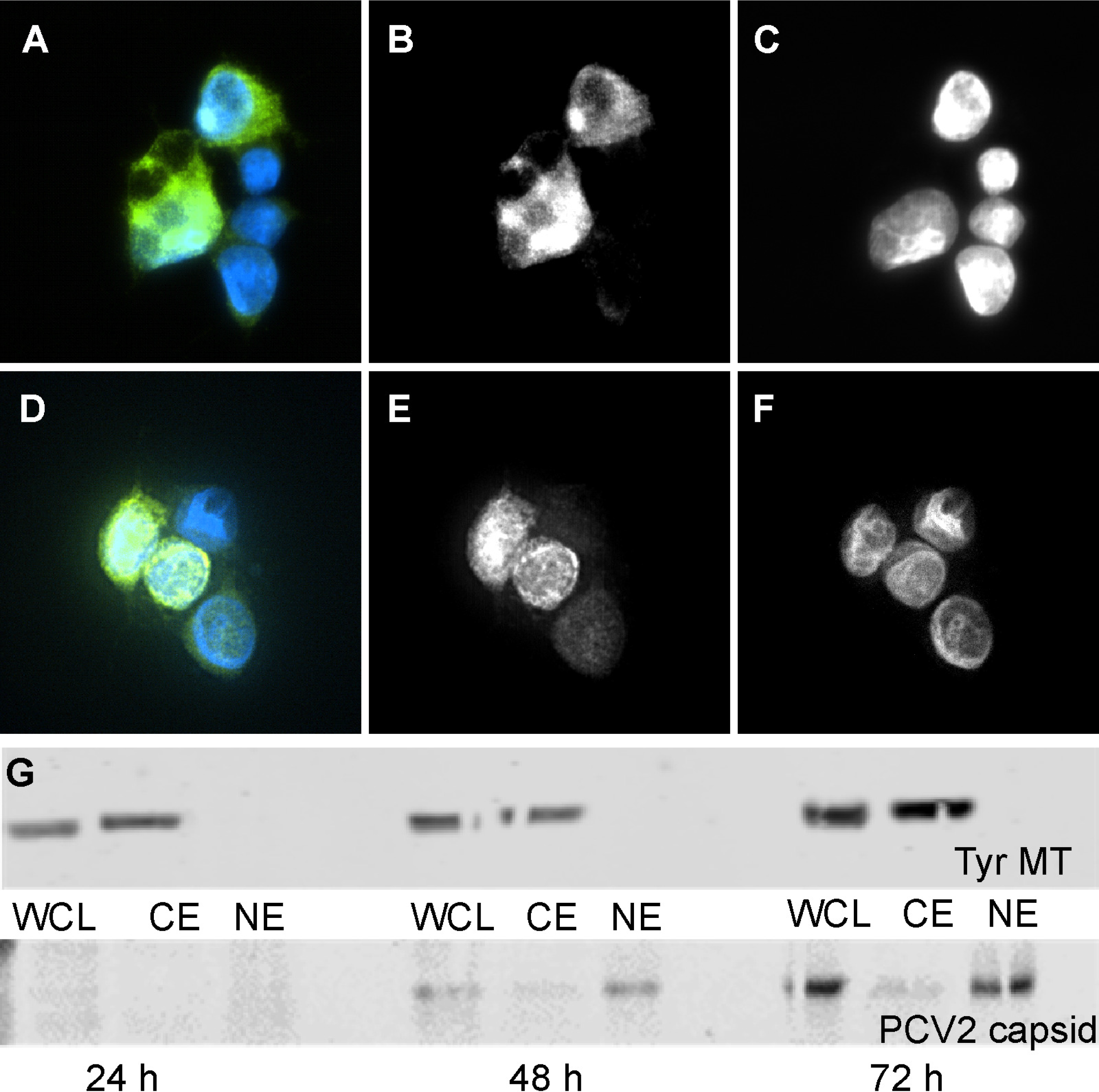
Location of expressed PCV2d capsid protein in transfected HEK293 cell. Transfected cells were fixed with acetone and labeled with mouse anti-PCV2 capsid monoclonal antibody to probe the capsid (green) and with DAPI (blue) to localize the nuclei. A) images from cells 48 hours post transfection with immunofluorescence from the capsid shown by the green channel and fluorescence from DAPI (nucleus labeling) by the blue channel. B) same image showing information from the green channel (capsid), C) same image showing information from the blue channel labeling (nucleus). D) images from cells 72 hours post transfection with immunofluorescence identical to A). E) same image showing information from the green channel (capsid), F) same image showing information from the blue channel (nucleus). G) Western blot analysis of whole cell lysate (WCL), cytoplasm (CE) and nuclear extracts (NE) collected 24, 48 and 72 hours post transfection. Top panel) The samples were probed for tyrosinated microtubules. Bottom panel) The samples were probed for PCV2 capsid protein. The presence of tyrosinated microtubules in CE but not in NE provided the quality control of cell fractionation.

Analysis of purified PCV2d VLP and its protein composition was confirmed by SDS-PAGE and Western blot analysis (Fig. 1B and C). Further examination by negative stained electron microscopy showed homogeneous spherical particles with smooth edges, slightly rough surface with a diameter of ~19 nm (Fig. 1D). The CP described in this study possesses the amino acid sequence identified from a number of recently isolated and reported PCV2d virus genome entries in GenBank, such as W233-12 isolated in Japan (BBE28610.1), CN-FJC011 isolated in China (AVZ66019.1), and England/15-P0222-09-14 isolated in England (ATN97185.1).

### Cryo-electron microscopy and image reconstruction of PCV2d VLP

We determined an icosahedral cryo-EM image reconstruction of purified PCV2d VLP to a resolution of 3.3 Å (Fig. 3A). The molecular envelope of the side chains allows us to confidently model the atomic coordinates of PCV2 (Fig 3B). The coordinates from the PCV2b crystal structure were manually fitted into the image reconstruction and appropriate modifications were performed to reflect the PCV2d amino acid sequence. Additional amino acids were built for the N-terminus and the fitted model was refined through several iterations of automatic refinement with Phenix suite and manual modeling using the program Coot (Fig. 3B). The final refinement statistics are shown in Table 1. We were able to model atomic coordinates from residue 36-231. Weak molecular envelopes preceding amino acid 36 and following amino acid 231 could be observed, but no models were built due to interpretation concerns. The PCV2 CP fold is the canonical viral jelly roll first visualized for the tobacco bushy stunt virus (11, 20). The jelly roll can also be described as a β-sandwich that is composed of two β-sheets. Each sheet consists of four strands (BIDG and CHEF) (Fig. 3C) and has a slightly right handed twist. The 60 CP pack together such that the β-sheets are normal to the *T=1* icosahedral particle. The loops connecting the strands BC, DE, FG and HI are 4 to 9 amino acids, while the loops connecting strands CD, EF and GH are 21 to 36 amino acids. The shorter loops define the surface of the capsid, while the longer loops are predominantly involved in CP-CP interaction. Given high sequence identity shared between the PCV2a, b and d sequences, it would be anticipated that no significant differences exist between their atomic coordinates. However, superimposing the subunits from these genotypes and generating root-mean standard deviation plots of equivalent Cα atoms proves otherwise (Fig 3C) (9, 21). The regions that exhibit significant diversity correspond to two of the surface-exposed loops consisting of amino acids 85-91 (section of loop CD), and 188-194 (section of loop GH) (Fig. 3C). The movement in loop CD and GH are coordinated, and a result of a single amino acid substitution at position 89 (loop CD) that is situated under loop GH. The larger amino acids of PCV2d (Leu89) and PCV2b (Arg89), as compared to PCV2a (Val89), push loop GH of PCV2d and PCV2b further away from loop CD. The Cα atom of PCV2a Thr189-Ser190 and PCV2b Thr189-Ala190 (loop GH) are ~1.4 and 1.3 Å apart, respectively. The Cα atoms for amino acids 189-190 in PCV2a and PCV2d are closer to one another, because of the smaller size differential between PCV2d Leu89 and PCV2a Val89. The change in the loops position may be a response to the immune system.

**Figure 3.**
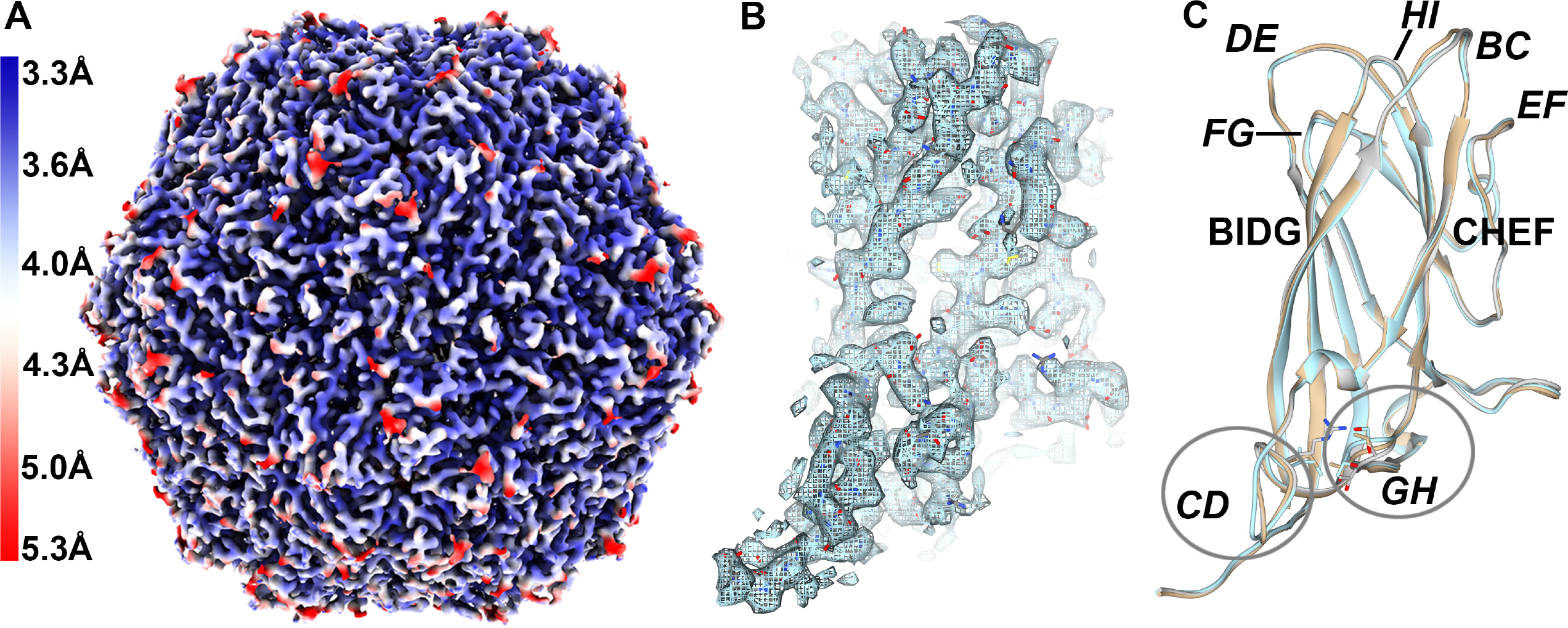
Structural study of the PCV2d VLP. A) Icosahedral cryo-EM image reconstruction of the purified PCV2d VLP colored according to the local resolution. The gradient color map on the left-hand side indicates the resolution for the colors. B) Extracted molecular envelope for a subunit demonstrates the quality of the image reconstruction, where amino acid side chains can clearly be seen. The atomic coordinates have been modeled into the image reconstruction. C) Structural overlay of the PCV2a (cyan), PCV2b (green), and PCV2d (salmon). Amino acids 89, 189 and 190 are shown as stick models. The β-strands (bold) loops (bold-italic) are labeled. Figures generated using UCSF Chimera and ChimeraX (57, 71).

**Table 1.**
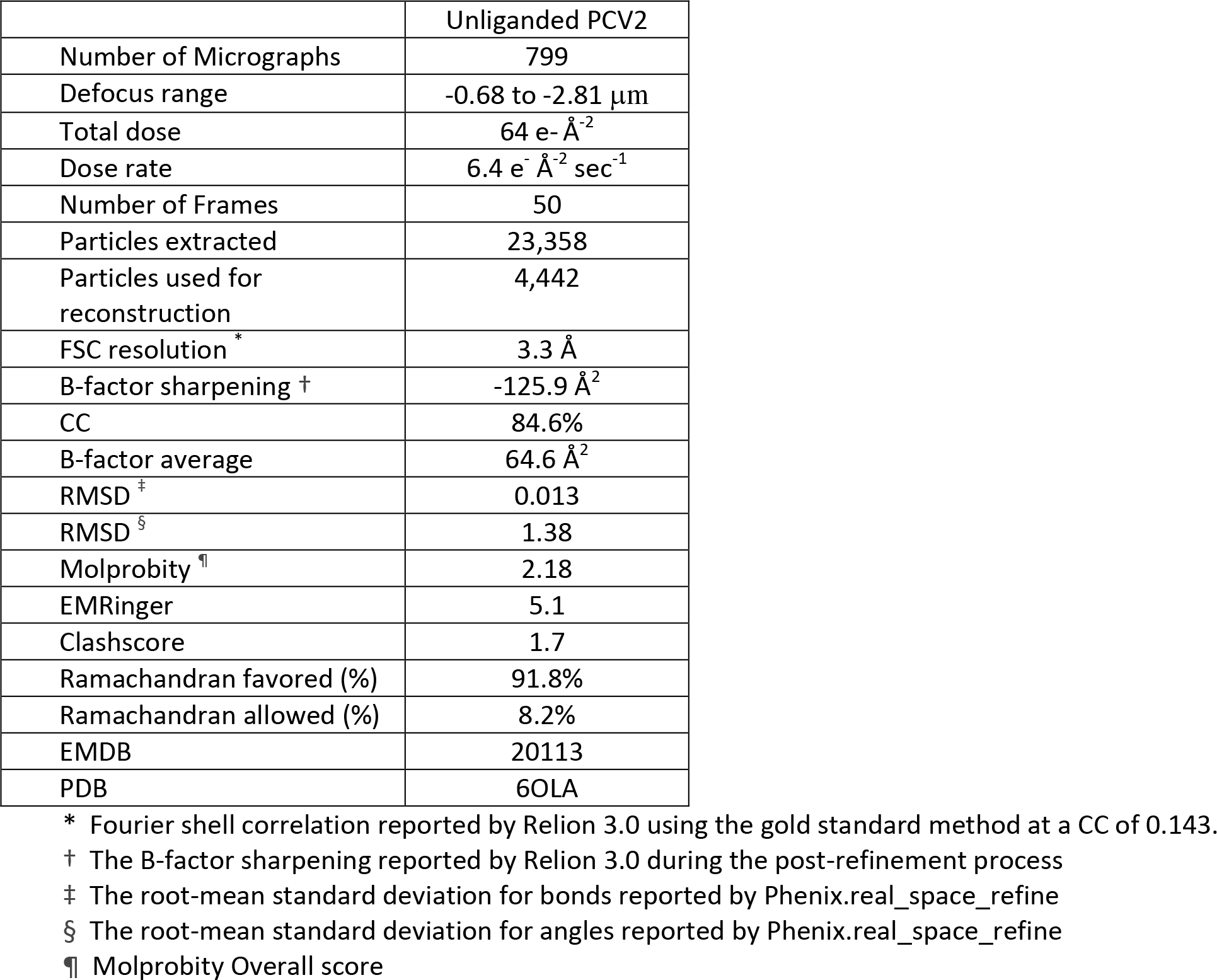

### The capsid packages cellular nucleic acid and recognizes a Pu-Pu-Py-Py nucleotide sequence

A near-central reconstruction slice was extracted from the cryo-EM image reconstruction and the density trace of pixel values was calculated in the horizontal and vertical directions (Fig 4A). Based upon the central slice of the reconstruction, the PCV2 VLP outer diameter is approximately 18.5 nm assuming a roughly spherical shape for calculation. The outer volume is therefore estimated to be 3.3×10^3^ nm^3^. The inner diameter using the same spherical shape estimation has a diameter of approximately 13 nm. Therefore, the inner region volume is approximately 1.2×10^3^ nm^3^. The radial profile of the image reconstruction indicates that a substantial amount of material is located within the capsid (Fig. 4A). We computationally measured the amount of material located inside the capsid by comparing the voxel count within the capsid shell to that inside the capsid -see methods and materials. A similar comparison performed for the asymmetric image reconstruction of the MS2 bacteriophage (EMD-8397) allowed us to calibrate the amount of material in our capsid to a sample with known RNA content. EMD-8397 is of sufficient quality to model the majority of the ssRNA genome (22). Our analysis indicates that in addition to the 60-copies of amino acids 1-42, which are believed to be in the interior of the capsid shell, there is scattering from an additional ~384 kDa of material. This is equivalent to ~1,200nt of ribonucleic acid. We also determined the RNA-protein ratio of PCV2d VLP using absorption light spectroscopy after correcting for light scattering (23).

**Figure 4.**
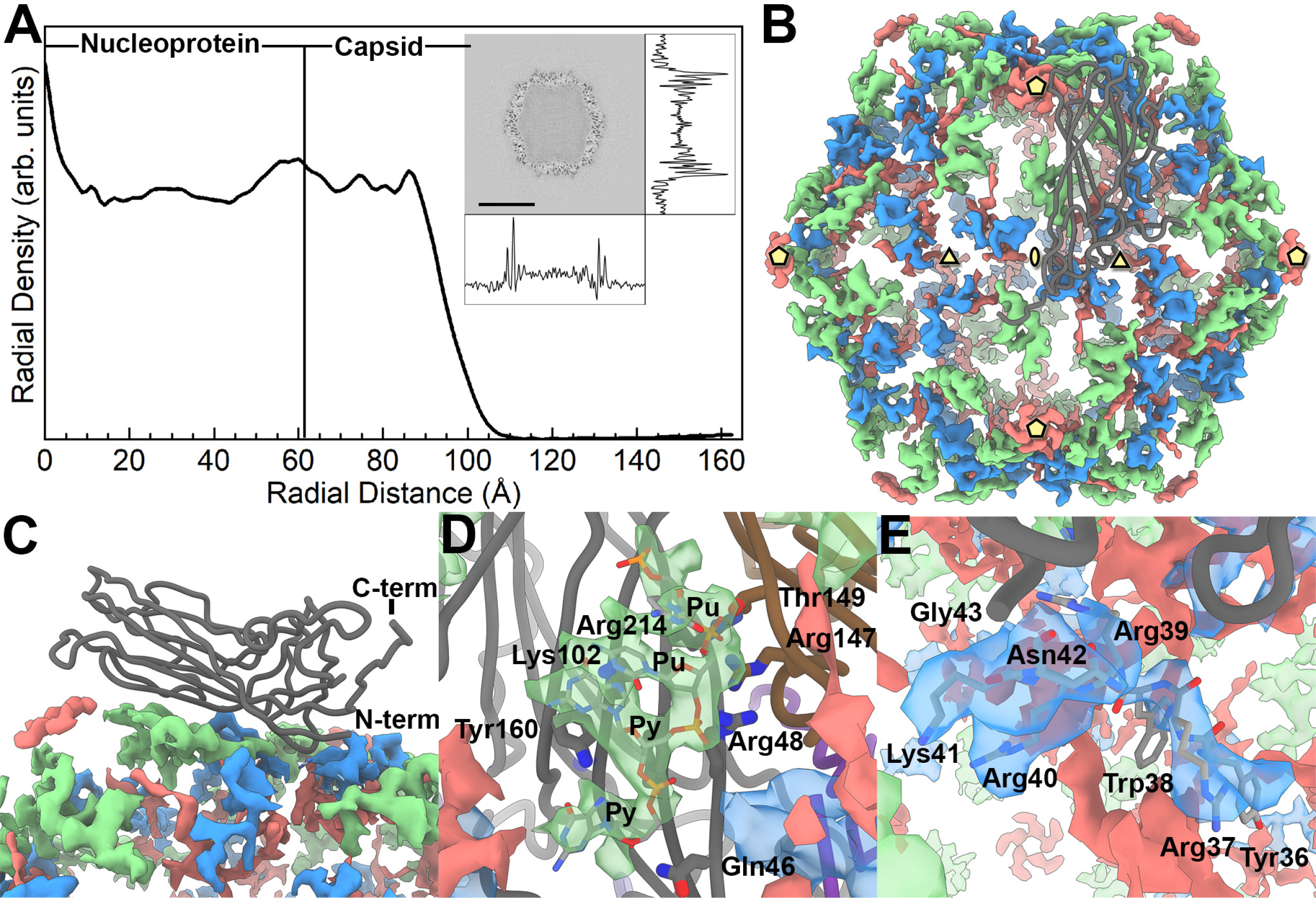
The inner content of the PCV2 capsid. A) A radial profile of the PCV2d cryo-EM image reconstruction. The capsid interior and shell are identified by assessing the cryo-EM image reconstruction. The inset is a central slice extracted from the cryo-EM image reconstruction, with the density trace of pixel values calculated in the horizontal and vertical directions. The radial profile demonstrates that number of voxels within the capsid is comparable to the capsid shell; thus, a substantial amount of material is located within the capsid interior. B) Strong difference peaks identified in the inner capsid. The icosahedral 5-, 3- and 2-fold axes of symmetry are identified by yellow pentagons, triangles and ellipses, respectively. A CP subunit is shown as a dark grey tube. We interpret the green colored difference peak to be a Pu-Pu-Py-Py tetranucleotide, the blue colored difference peak to be amino acids 36-41 of the PCV2 N-terminus, and the red colored difference peak to be “unidentified”. C) Side view showing the CP subunit and the difference peaks. D) Close up of the tetranucleotide that has been modeled into the difference peak (green) located near the 3-fold axes of symmetry, and the CP amino acids in proximity. Gln46 (strand B), Arg48 (strand B), Lys102 (strand D), and Arg214 (strand I) of one subunit, and Thr149 (strand F) and Arg147 (strand F) from a neighboring subunit form hydrogen bonds and electrostatic interaction with the phosphate backbones of the tetranucleotide. Tyr160 (strand I) forms π-bond overlap with the first Py in the tetranucleotide. E) Close up of the PCV2 N-terminus modeled into the difference peak (blue) located near the 3-fold axes of symmetry. Amino acids 36-42 are labeled.

We then used adsorption spectroscopy to measure the content of the purified PCV2d VLP following the protocol described by Porterfield and Zlotnick (23). The measurements allow one to subtract scattering (absorbances at 360nm and 340nm) from the sample and calculate the amount of RNA (absorbance at 260nm) present as a ratio of the amount of protein (absorbance at 280nm) present. Five measurements were made to yield an average of 23.8 nucleotides per CP. This is equivalent to 1,428 nucleotides per capsid, and in agreement with our estimations using the cryo-EM image reconstruction of PCV2d.

To interpret the inner composition of the PCV2d VLP image reconstruction, we generated a molecular envelope (difference map) representing the inner composition of the capsid by subtracting a 3.3 Å molecular envelope calculated from our coordinates (amino acids 43-231) from our 3.3 Å cryo-EM image reconstruction (Fig 4B, C). Strong difference peaks are observed near the icosahedral 3-fold and between the 5- and 2-fold axes of symmetry. We model a short oligonucleotide (Pu-Pu-Py-Py) into the difference peak between the 5- and 2-fold axes of symmetry. The backbone ribose and phosphates of the oligonucleotide form hydrogen bonds and electrostatic interactions with Gln46, Arg48, Lys102 and Arg214 of one subunit, and Arg147 and Thr149 of a neighboring subunit. The side chains from Arg48, Arg147 and Arg214 stack like a zipper to form a positive patch on the inner surface of the capsid. The charge on the patch is neutralized by the two phosphates in the RNA backbone. The second Pu of the oligonucleotide forms π-interactions with Tyr160 of the PCV2d CP (Fig 4D). Two observations support our model: the modeled coordinates demonstrate the nucleotide bases to stack and form π-bond overlap, and the coordinates of these nucleotides overlay with the coordinates of nucleotides that were modeled into the beak and feather disease virus (**BFDV**) crystal structure when the PCV2 and BFDV CP are superposed -BFDV is a member of the *Circoviridae* family and its CP shares 32% sequence identity with the PCV2 CP (24).

### The PCV2 N-terminus is in the interior of the capsid and located near the icosahedral 3-fold axis of symmetry

The N-terminus of *Circovirus* capsid proteins are highly charged and predicted to be an intrinsically disordered region -a region with no secondary structure (25). We recently provided experimental support for this prediction by measuring the circular dichroism spectrum of the PCV2b N-terminus peptide (amino acids 1-42) (Manuscript under review). The crystal structure and cryo-EM image reconstruction of the PCV2b VLP and the cryo-EM image reconstruction of the PCV2a VLP both strongly suggested that the N-terminus is located inside the capsid (9, 11, 26). Half of the amino acids in the N-terminus are composed of Arg or Lys amino acids; thus, its high positive charge and potential location suggests that it may be capable of interacting with nucleic acid packaged in the capsid. The difference peak near the 3-fold axes of symmetry are adjacent to the modeled N-terminus of PCV2 (amino acid 43), and we have modeled six amino acids into this difference peak to extend the N-terminus to amino acid 36 (Fig. 4E). The N-terminus begins as a single turn helix that descends into the inner portion of the capsid. Additional molecular envelop for the N-terminus can be observed; however, we refrain from modeling this section as we cannot confidently interpret the direction of the polypeptide chain and identify the amino acids by their side chain molecular envelope. The reduced quality of the molecular envelope in this region is most likely due to the N-terminus not adopting the icosahedral symmetry of the capsid. Unfortunately, symmetry expansion, signal subtraction, and focused classification of this region was unable to improve the quality of the molecular envelope. Consequently, we do not know the location of amino acids 1-35 of the CP. A recent manuscript by Mo *et al.* models the N-terminus of PCV2 near the icosahedral 5-fold axes of symmetry (19). We will discuss the discrepancy between the two models in the discussion section.

Difference maps calculated from the PCV2a and PCV2b image reconstructions display the tetranucleotide molecular envelope located between the icosahedral 2- and 5-fold axes of symmetry to be conserved (Fig 5A-C). Surprisingly, the molecular envelope for the PCV2a N-terminus is rotated ~90° with respect to its equivalents in the PCV2b and PCV2d image reconstructions (Fig 5A-C). Radial plots of the image reconstructions suggest that the inner contents of the PCV2a image reconstruction consist of the least amount of packaged material (Fig 5D). Perhaps the distinct positioning of the N-terminus in the PCV2a image reconstruction is a result of the lesser amount of packaged material.

**Figure 5.**
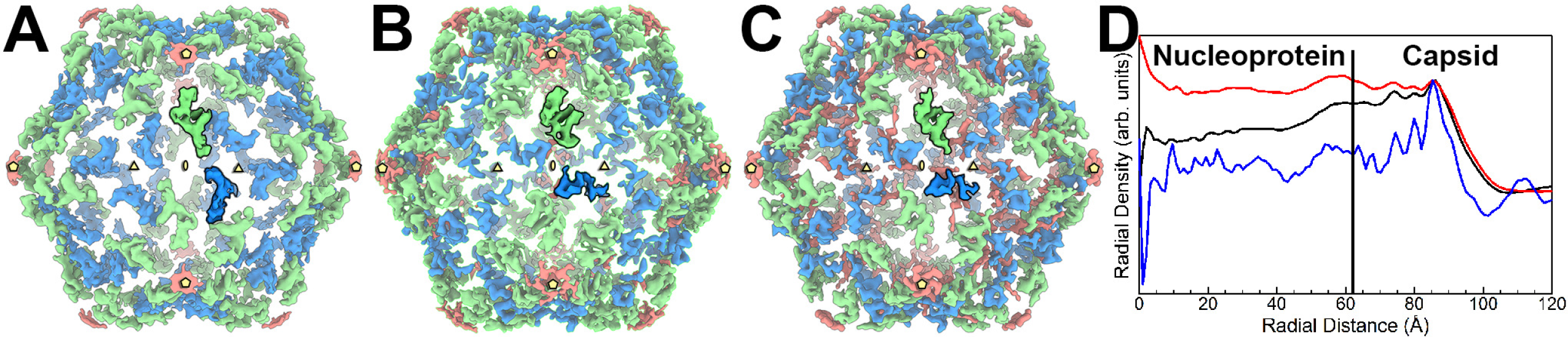
Inner capsid difference peaks of PCV2a, PCV2b and PCV2d. Comparing the difference peaks in the inner capsid of PCV2a (A), PCV2b (B) and PCV2d (C) reveals a conserved location for the tetranucleotide molecular envelope (green) and differences for the N-termini molecular envelope (blue). The orientation of the PCV2b and PCV2d N-termini are conserved; however, the N-termini of the PCV2a are rotated ~90° clockwise. The icosahedral symmetry elements are shown using the same convention as Fig 4B. The molecular envelopes for a single subunit are painted with bold colors. D) Radial plots of PCV2a (blue), PCV2b (black), and PCV2d (red). The plots are normalized and scaled to one another to simplify the comparison. The PCV2d image reconstruction possesses the greatest amount of content within its capsid, while the PCV2a image reconstruction possesses the least amount of content within its capsid.

### PCV2a, b and d genotypes demonstrate different diversity and variability

Several phylogenetic studies have compared the amino acid sequence of PCV2 CPs to differentiate PCV2 into genotypes (6, 7, 14). Indeed, Franzo and Segales recently identified eight PCV2 genotypes (a-h) (14). A report by Wang *et al.* mapped the sequence variation in 50 PCV2 CP sequences onto the crystal structure and identified regions of variability (8); however, since then a significant number of CP sequences have been deposited into GenBank that requires revisiting the analysis. We asked, do amino acids that exhibit the greatest diversity (lowest sequence identity) or greatest variability (AL2CO, entropy) cluster on the 3D structure of the CP, and is the clustering conserved among PCV2a, b, and d genotypes (27). We chose to concentrate on these three genotypes because the larger number of GenBank entries allow us to reach a statistically meaningful conclusion. We initiated our analysis by downloading the nucleotide sequences used by Franzo and Segales in their study (PCV2a: 675; PCV2b: 1984; PCV2d: 1491) from GenBank (14). We note that sequences arising from recombination have been removed by Franzo and Segales. We then converted the nucleic acid sequences to amino acid sequences using the NCBI ORFfinder program (28) and removed all redundant amino acid sequences from each genotype using the CD-HIT server (29). A total of 317 PCV2a, 501 PCV2b and 368 PCV2d unique sequences were aligned for each genotype and were mapped onto the PCV2a, PCVb and PCV2d atomic coordinates (Fig 6). The PCV2a genotype demonstrates the greatest sequence diversity with the lowest sequence variability -the two most diverse sequences share 86.3% identity (AF364094.1 vs. AIQ85146.1) (Fig 3C and 6A). The most diverse positions are in loops BC, CD, EF, GH, and HI while the most variable positions are in loop CD. The high sequence diversity in the apposed loops BC and HI is suggestive of a conformational epitope (*conf1*). Similarly, the high sequence diversity in the apposed loops CD and GH is suggestive of a conformational epitope (*conf2*). Indeed, Shang *et al.* demonstrated that PCV2 neutralizing antibodies 2B1, 7F4, and 6H9 bound to PCV2 isolates NB0301, SX0201, HZ0301, HZ0201, TZ0601, JH0602, but not ISU-31 (30). The difference in sequence between ISU-31 and the remaining PCV2 isolates maps to loops CD, GH and HI. Consequently, antibodies 2B1, 7F4, and 6H9 may bind to either *conf1* or *conf2*. The PCV2b genotype exhibits a lesser sequence diversity, but a greater sequence variability –the two most diverse sequences share 84.5% identity (ABL07437.1 vs. ABR14586.1) (Fig 3C and 6B). The most diverse positions are in loop BC and the most variable positions are in loops BC, CD, DE, and HI. The PCV2d genotype exhibit distinct sequence diversity and variability when compared to the PCV2a and b genotypes -the two most diverse sequences share 83.3% identity (KP824717.1 vs. AVR51420.1). For example, the PCV2b genotype exhibits greater sequence diversity in loop BC and variability in loop CD, while PCV2d exhibits greater sequence diversity in loop GH and diversity in loop EF (Fig 3C and 6C). Not shown in Fig 4 are the last few C-terminal amino acids of the CP, as none of the structural studies have been successful in visualizing these amino acids. These amino acids are highly variable within each genotype, and between genotypes (2, 5, 7, 14). PCV2b and PCV2d also exhibit greater sequence variability in the inner surface of the capsid when compared to PCV2a (Fig. 6A-C).

**Figure 6.**
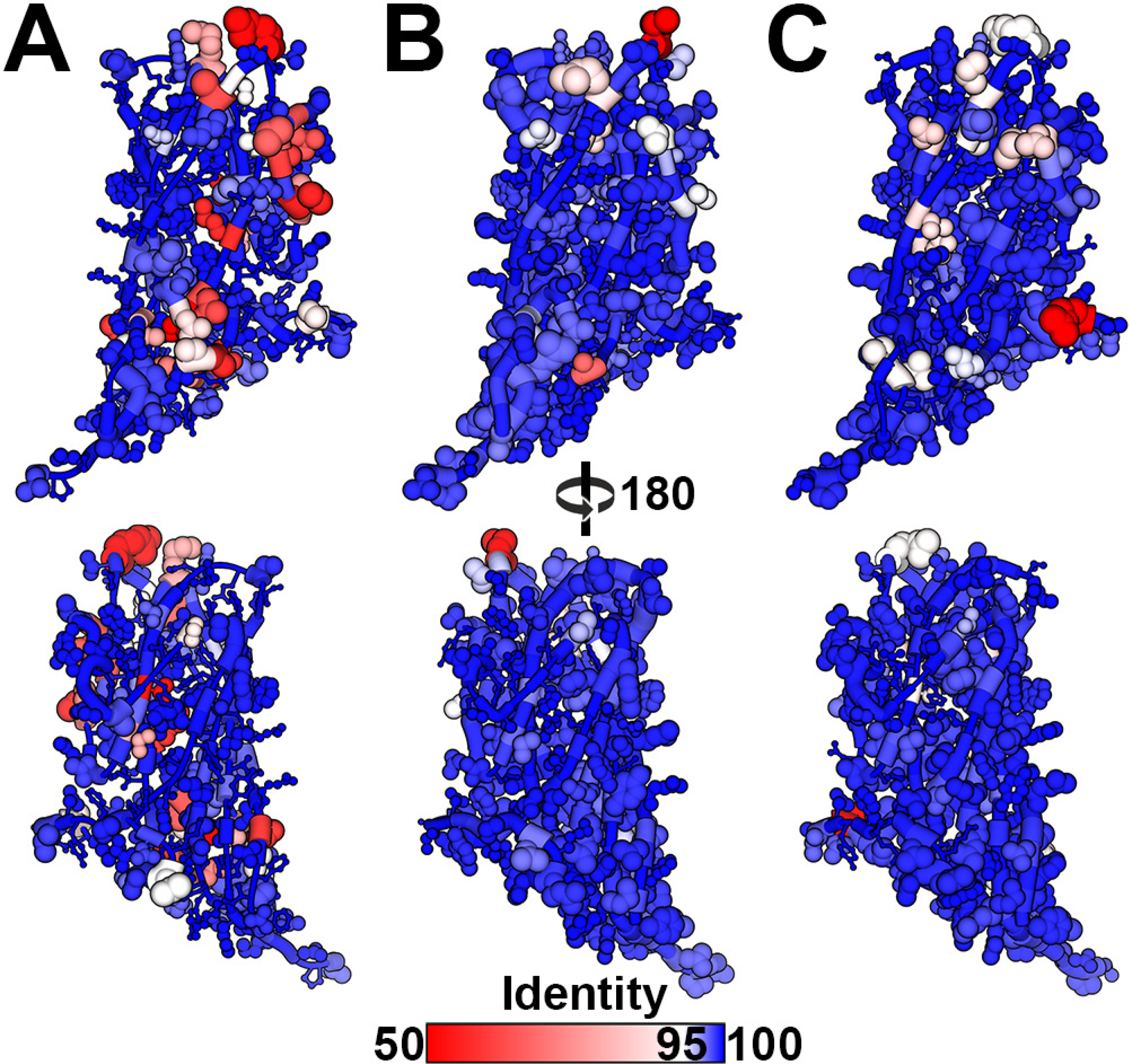
Sequence diversity and variability in the PCV2a, b, d genotypes. A) Amino acid sequence alignment information plotted on the PCV2a, b, and d coordinates. The red-to-blue color gradient represents diversity (sequence identity) -gradient shown at the bottom of the image. The size of the atoms and tube represent variation (sequence entropy), with smaller atoms/tubes representing lower entropy and larger atoms/tubes representing greater entropy. Lower entropy indicates fewer amino acids present in the alignment at a position, and larger entropy indicates more amino acids present at a position. The plotting of variation allows one to appreciate the frequency of different amino acids at each position. Top, amino acids facing the capsid exterior. Bottom, amino acids facing the capsid interior. A) Attained from 317 unique CP entries plotted on the surface of the PCV2a atomic coordinates (PDB entry 3JCI). B) attained from 501 unique CP entries plotted on the surface of the PCV2b atomic coordinates (PDB entry 6DZU). C) attained from 368 unique CP entries plotted on the surface of the PCV2d atomic coordinates (PDB entry 6OLA).

### Highly conserved amino acids form patches on the capsid surface

The sequences discussed above were expanded to include all PCV2 GenBank entries, then reduced to a unique set of amino acid sequences (1,278 entries). The sequence alignment was used to generate a WebLogo diagram to observe the sequence conservation among all PCV2 genotypes (Fig. 7A). The sequences with the greatest diversity (GenBank entries: AVZ65995.1 and ALK04312.1) share 74.4% sequence identity. While the WebLogo diagram successfully demonstrates the frequency of occurrence for the most popular amino acid(s) at each position, it dampens the frequency of occurrence for the less popular amino acids. Indeed, substitutions are observed within the hydrophobic core of the protein, at the intersubunit interface, and amino acids on the outer and inner surface of the capsid. Moreover, only Met1, Pro15 and Arg147 are absolutely conserved. The analysis suggests that the capsid is capable of tolerating mutations at nearly every position while remaining an infectious virus. To visualize the sequence conservation at each amino acid position, we plotted the sequence alignment information onto the PCV2d atomic coordinates using the ConSurf server (Fig. 7B) (31). The figure displays a significant degree of nonconserved substitutions. Regions colored in red experience the greatest degree of change while regions in blue experience the least degree of change. Highly conserved amino acids are peppered across the structure; however, two sets of amino acids form patches on the surface of the capsid. These include Tyr55, Thr56, Met71 and Arg73, and Pro82, Thr170, Gln188, Thr189 and Val193 (Fig. 7C). The presence of these patches suggests that they may play important biological roles in the PCV2 life-cycle (e.g. cellular interaction, capsid dynamics/stability). Given the conservation of these amino acids, it would be interesting to test if vaccines capable of inducing antibody production to these patches (conformational epitopes) are able to neutralize all PCV2 genotypes (Fig. 7C).

**Figure 7.**
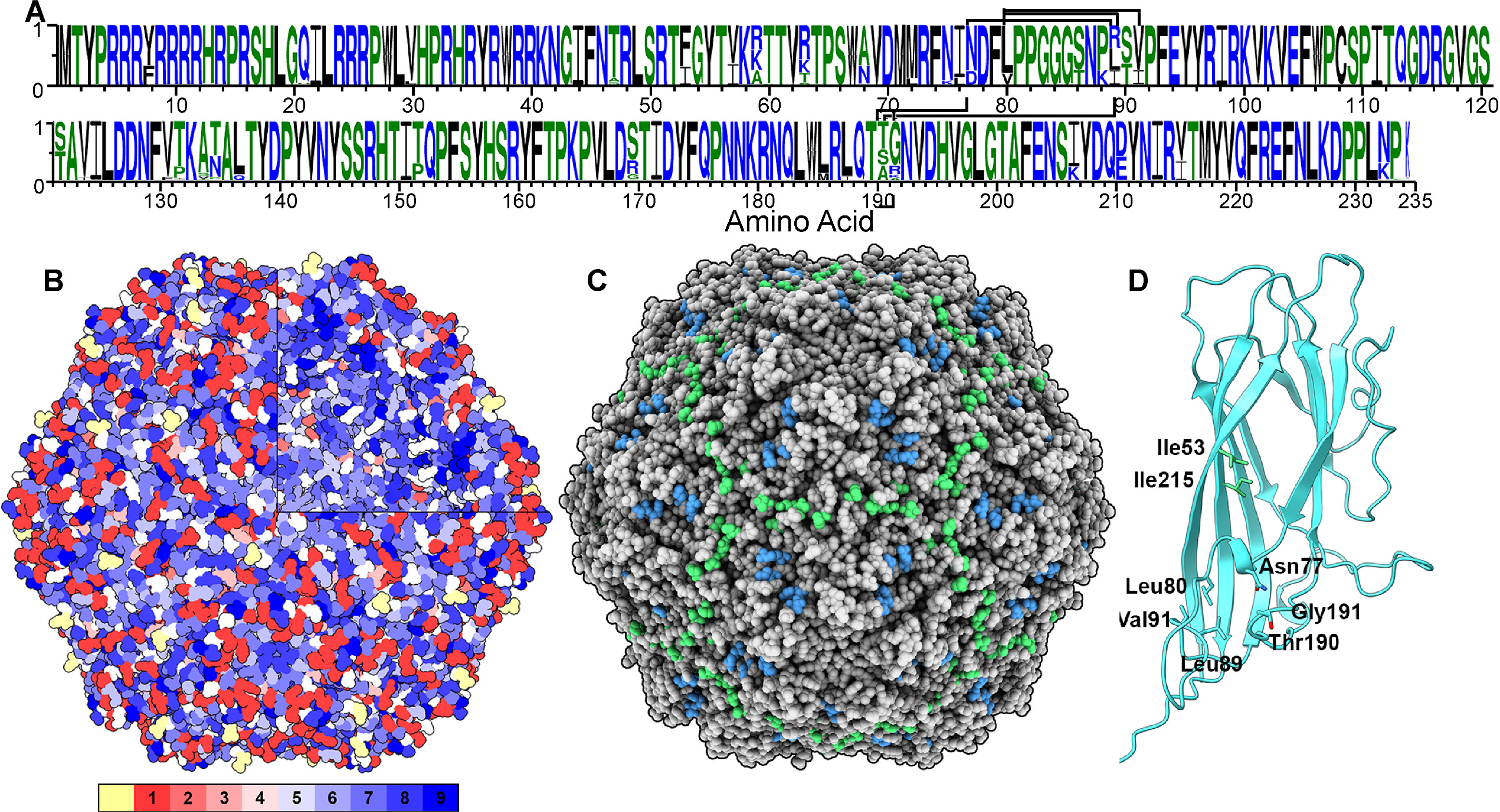
Sequence comparison of 1,278 PCV2 capsid protein entries plotted on the PCV2d atomic coordinates. A) The sequence alignment by the Clustal Omega server was used to generate the WebLogo diagram to demonstrate sequence variation. The horizontal axis of the alignment indicates the amino acid and the vertical axis indicates its observed frequency. Bars connecting amino acids 77, 80, 99, 91, 190, 191 (black), and 53, 215 (grey) represent the evolutionary coupled clusters shown in panel D. B) Space filling model of the PCV2d atomic coordinates with a modified color-coding scheme of ConSurf. The color bar at the bottom indicates the degree of conservation determined by the ConSurf server. The yellow box indicates insufficient data as determined by the server (1 indicates poorly conserved and 9 indicates highly conserved mutations). The top right quadrant of the VLP surface has been removed to display the sequence conservation in the interior of the capsid. Image made with UCSF ChimeraX and colored using flat lighting. C) Highly conserved amino acid on the capsid surface (*conf1:* amino acids 82,170,188,189 and 193 in green, *conf2:* amino acids 55, 56, 51 and 73 in blue) form patches. Antibodies directed against these patches (conformational epitopes) may exhibit broadly neutralizing capability. D) Ribbon cartoon of a PCV2d subunit. Amino acids in stick are evolutionary coupled together, as determined using the plmc. MATLAB 2019, and EVzoom programs. Figures generated using UCSF ChimeraX (71).

### Evolutionary coupled mutations differentiate the PCV2 genotypes

The large number of CP sequences allowed us to ask if any of the amino acid positions are evolutionary constrained (32). For evolutionary constrained amino acids, mutation of one amino acid requires mutation of the other amino acids. In the simplest of cases this may be because the amino acids pack against one another in the structure of the protein, such that mutation to a larger amino acid in one position requires mutation to a smaller amino acid in the second position for proper packing to occur. Such information can be used to predict the fold of a protein or identify functionally important sites (32–34). Evolutionary coupling (**EC**) measurements determined from the 1,278 unique sequences indicates that two independent locations in the structure demonstrate coupling. The first location consists of amino acids 53 and 215, and the second location consists of amino acids 77, 80, 89, 90, 190 and 191 (Fig. 7C). Sequence differences in the second location are responsible for the structural diversity observed between PCV2a and PCV2b/d in loops CD and GH (Fig. 3C) (9, 21). The EC results may indicate that the coupled amino acids are functionally relevant. Solvent accessible surface area calculations with the GetArea server (http://curie.utmb.edu/getarea.html) indicates that amino acids 77, 80, 89, and 90 are more than 75% buried, while amino acids 190 and 191 are exposed to the solvent (35). Amino acids 190 and 191 help define a neutralizing epitope on the surface of the capsid -see below.

### Sequence variation on the capsid surface may be a response to neutralizing antibodies

We used the atomic coordinates of PCV2d to identify amino acids on the surface of the VLP with side chains exposed to solvent (53, 55-56, 58-64, 70-71, 73, 75, 77-78, 82-83, 85, 88-89, 102, 113, 115, 123, 127, 131-137, 148, 155, 156, 158, 161, 166, 168-170, 188-191, 194, 204, 206-208, 210, 229-234). These amino acids may be antigenic determinants, as the side chains provide a surface for antibody interaction. Continuous sequences encompassing the above amino acids are shown in Fig. 8A-H, with the most variable amino acids highlighted (53, 57, 59, 60, 63, 75, 77, 88, 89, 131, 133, 134, 136, 169, 190, 191, 211, 215, 232). Mahe *et al.* used PEPSCAN analysis of PCV1 and PCV2 CP to identify linear epitopes of CP (amino acids 65-87, 113-139, and 193-207) capable of interacting with PCV2 antisera (13). Khayat *et al.* used this information to identify the amino acids of the PCV2 CP crystal structure (amino acids 70, 71, 77, 78, 113, 115, 127, 170, 206 and 207) that may be responsible for the observed PEPSCAN reactivity (11). Antigenic subtyping experiments by Lefebvre *et al.* and Saha *et al.*, where panels of neutralizing monoclonal antibodies are tested for their ability to bind to different strains of PCV2, demonstrated that amino acid positions 59, 63, 88, 89, 130, 133, 206 and 210 are responsible for differentiating antibody binding (36, 37). Huang *et al.* demonstrated that amino acid 59 resides in a conformational epitope for the neutralizing antibody 8E4 (38). Franzo *et. al*. described the evolution of PCV2 before and after the introduction of vaccination to involve changes in amino acids 59, 191, 206, 210, 228 and 232 (39). Except for amino acid 130, which is buried in a subunit-subunit interface, these amino acids are on the surface of the capsid and undergo extensive substitutions. It is certainly possible that a combination of two or more proximal regions may help define conformational epitopes (e.g. Fig 8A, D, G; Fig 8E-F; Fig 8B, D; Fig 8 D, H; Fig 8C, H; Fig 8C, F).

**Figure 8.**
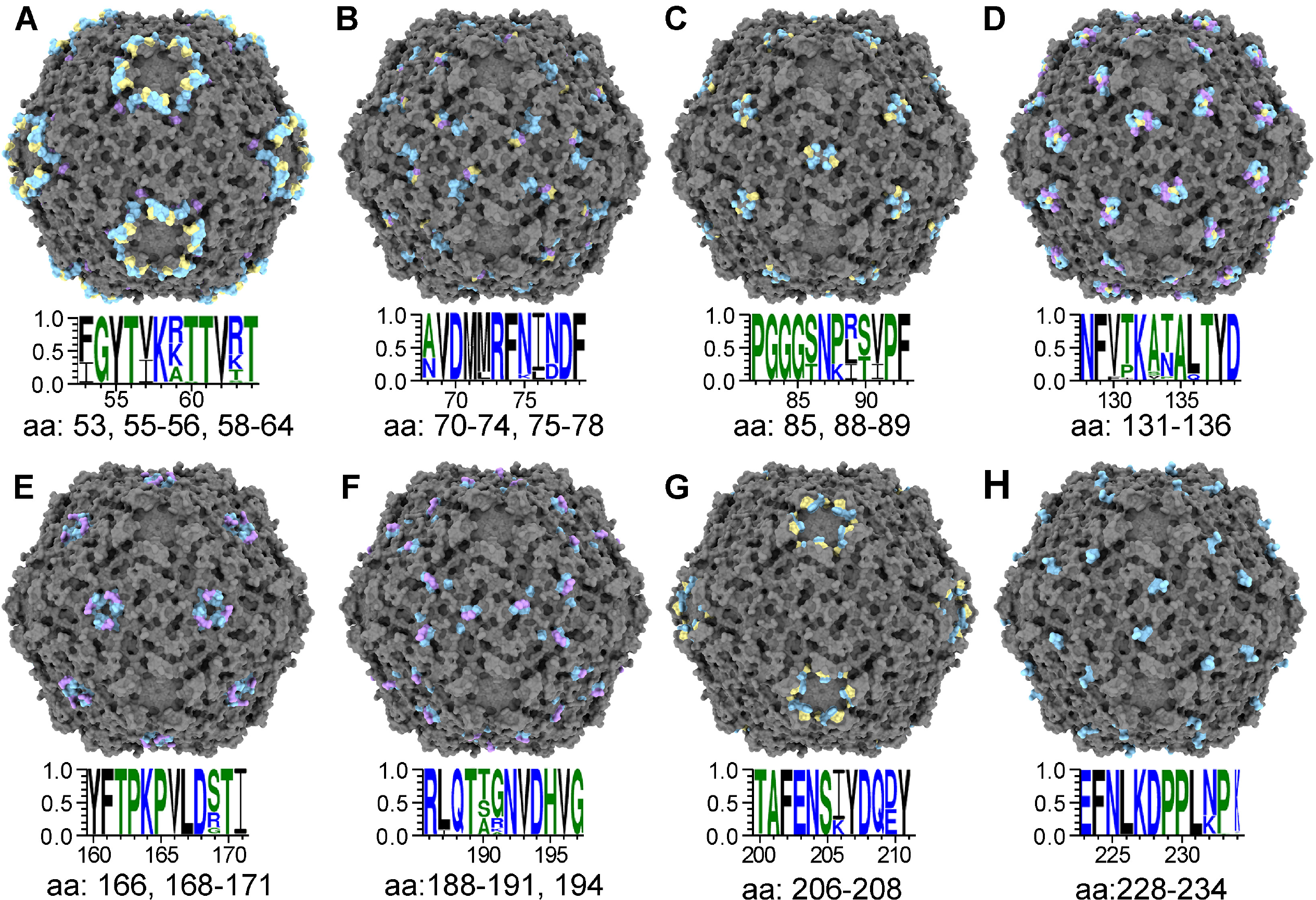
Sites of antibody neutralization. Top) Space filling model of the PCV2d atomic coordinates with the surface exposed amino acids colored in cyan, sequence variable amino acids in purple, and antibody binding amino acids in yellow. Middle) WebLogo diagram of 11 amino acids containing the solvent exposed amino acids. Bottom) Amino acids on the surface of the capsid. The image concentrates on continuous amino acids, but the proximity of two such regions can define a conformational epitope.

## Discussion

Porcine circovirus 2 genome encodes for four known proteins: a replicase (ORF1) responsible for genome replication, a capsid protein (ORF2) responsible for generating the capsid shell, and an ORF3 and ORF4 that are believed to regulate cellular apoptosis (40, 41). PCV2 demonstrates the ability to rapidly mutate and evolve into novel genotypes -an ability that is governed by the accumulation of point mutations (42, 43) and genome recombination (44). Phylogenetic analysis of the PCV2 CP sequence indicates that eight genotypes are distributed globally (PCV2a-h) (14). The larger deposition of entries for PCV2b in GenBank suggests that it is the dominant genotype; however, the recent decrease in PCV2b and increase in PCV2d depositions suggests that there may be a shift from PCV2b to PCV2d (5). Given the history of shifts in PCV2 genotypes, it would not be too surprising if additional shifts occurred. To understand why the genotype shifts are occurring, it is necessary to establish an expression and purification protocol to rapidly acquire large quantities of PCV2 VLP for structural and biophysical studies. It is important to structurally characterize the PCV2 capsid as it is the entity under selective pressure by the environment (e.g. immune system and cellular interaction), and sequence information sometimes is not enough to distinguish differences between genotypes. Consequently, we have established a rapid and robust mammalian expression system for producing large quantity of PCV2 VLP. The study described in this paper is the first system where mammalian cells have been utilized to generate large quantity of VLP. The relatively easy and rapid protocol of the mammalian expression system provides a tremendous advantage to the timely and involved protocol necessary for baculovirus generation. The mammalian expression system described here can generate VLP four days from when the expression plasmid is attained, whereas the fastest baculovirus expression protocol can generate VLP in eight days (45). The decrease in time necessary to generate VLP is particularly advantageous when mutagenesis experiments are to be performed. Our expression protocol can also be used to generate capsids assembled as a multivalent mosaic that simultaneously displays neutralizing epitopes of several genotypes. Immunofluorescence and Western blot analysis of CP expression revealed that CP can be detected in the cytoplasm at 24 hours, but only detect in the nucleus at 72 hours. This suggests that the VLP is unable to exit from the nucleus of mammalian cells within the first 72 hours of expression.

The molecular envelope of the 3.3 Å PCV2d cryo-EM image reconstruction allows us to confidently model amino acids 36 to 231, and a Pu-Pu-Py-Py tetranucleotide. Comparison of the PCV2a, b and d atomic coordinates attained from cryo-EM studies identifies differences in the conformations of the surface-exposed loops CD (amino acids 86-91) and GH (amino acids 188-194) (Fig. 3C). The movement of these loops are coupled to one another and thus suggestive that the difference is not happenstance. The difference between PCV2a and PCV2b in the positions of these loops provides a structural description for the genotype shift observed in 2003 from PCV2a to PCV2b (15, 16). Such a difference could not have been identified by sequence analysis alone. Indeed, the structures indicate that substitution at a single position (amino acid 89) is responsible for the difference in loop positions. The movement in loop position may be a response to neutralizing antibodies -see below. The Pu-Pu-Py-Py tetranucleotide coordinates modeled into our PCV2 image reconstruction is the first observation of ordered nucleic acid in a PCV2 capsid. The nucleic acid is most likely to be RNA as VLP assembly occurs in the nucleus of the mammalian expression system. Hydrogen bond and electrostatic Interactions are observed between Gln46, Arg48, Lys102 and Arg214 of one subunit, and Arg147 and Thr149 of a neighboring subunit with the backbone of the tetranucleotide. The base from the second Pu of the tetranucleotide forms π-interactions with the side chain of Tyr160. All four bases of the tetranucleotide make π-interactions. Interestingly, the placement of the coordinates with respect to the CP is nearly identical to that of the beak feather disease virus - another member of the *Circoviridae* family (24). The position for some of the amino acids that interact with the PCV2 nucleic acid overlaps with the position of amino acids that interact with the BFDV ssDNA, while others are unique to PCV2 or BFDV. PCV2 and BFDV share 32% sequence identity.

Amino acids 36-42 modeled into the N-terminus of the PCV2d image reconstruction are located inside the PCV2 capsid near the icosahedral 3-fold axes of symmetry (Fig 4D). Amino acids 43-50 form the B β-strand of the viral jelly-roll and remain in the interior of the capsid until amino acid 51, which is exposed on the surface of the capsid (11, 26). Interestingly, there is a discrepancy between our model and that recently described by Mo *et al.* (19). Mo *et al.* model amino acids 33-42 to interact with amino acids 17-27 near the icosahedral 5-fold axes of symmetry. Moreover, Mo *et al.* model amino acids 43-49 to form a loop that points to the exterior of the capsid near the 3-fold axes of symmetry. Unfortunately, a direct comparison between our model and that of Mo *et al.* cannot be made because the coordinates for their N-terminus model has not been deposited into the PDB or EMDB (19). The different models may be a result of the different expression and assembly systems used by these studies; we use a mammalian expression system where the VLP assemble in the nucleus of the cell whereas Mo *et al.* use an *E. coli.* expression system with *in vitro* VLP assembly. We note that our model is also consistent with the cryo-EM image reconstructions of PCV2a and PCV2b VLP attained from insect cell expression systems -where amino acids 43-50 are modeled in the interior rather than exterior of the capsid (9, 26). Further studies are necessary to clarify the discrepancies between the two models.

Comparison of difference maps calculated from subtracting the cryo-EM image reconstructions of PCV2a, b, and d from their respective atomic coordinates (amino acids 42-231) reveals similarities and differences between the interior of PCV2a and PCV2b/d (Fig 5A-C). All difference maps identify similar molecular volumes describing the tetranucleotide we’ve modeled into the PCV2d image reconstruction. However, the positions of the molecular envelopes for the PCV2a N-termini are different than those of PCV2b/d. While the N-termini for the three genotypes locate near the intra-facet icosahedral 3-fold axes of symmetry, the PCV2a molecular envelope is rotated 90° clockwise such that it points away from the 3-fold axes of symmetry (Fig 5A-C). Possibly, the position of the PCV2a N-terminus is correlated to the lesser amount of material present inside the PCV2a capsid (Fig 5D). If so, the N-terminus may act as a sensing switch for regulating capsid assembly. We are exploring this possibility with biochemical and structural studies.

Phylogenetic analysis of the PCV2 CP sequences has recently been used to identify eight genotypes (a-h) (14). We mapped the diversity and variability present in the CP sequences of the PCV2a, b, and d genotypes onto their respective atomic coordinates (Fig 6A-C) (9, 26). The plots, coupled with published work, suggest two conformational epitopes on the capsid surface. The plots also demonstrate that PCV2a exhibits the greatest sequence diversity but lowest sequence identity (Fig 6A). Both PCV2b and d exhibit high sequence variability (Fig 6B and 6D). The increased variability in the PCV2 genotypes may be a result of the selective pressure induced by the vaccination program in 2006 (46). The number of CP sequence entries into GenBank was dominated by PCV2a prior to the introduction of the program but was quickly shadowed by PCV2b entries. The number of PCV2d entries is greater than PCV2b entries for 2016 (14). We note that the sequence diversity observed in these three genotypes are sufficiently high such that a single sequence is not truly representative of the genotype when one considers antibody escape mutations. The diversity in CP sequences suggests that PCV2 may indeed be regarded as viral quasispecies (47).

Alignment of 1,278 unique PCV2 CP sequences indicates that only three amino acids (Met1, Pro15 and Arg147) are absolutely conserved, and that the remaining positions in the sequence have undergone mutations (Fig 7B). This demonstrates the remarkable plasticity of the capsid structure to undergo mutation while maintaining an infectious virus, and thus highlights the capacity of PCV2 to respond to natural selection. The sequence analysis coupled with the structural information identify conserved sites on the capsid surface (*conf1* and *conf2*) that may be taken advantage of for designing vaccines capable of eliciting broadly neutralizing antibodies (Fig 7C). Evolutionary coupling analysis of the sequence alignment identified two regions that underwent co-evolution. One of these regions correlates with the structural diversity observed between PCV2a and PCV2b/d in loops CD and GH (Fig 7D). The continuous regions on the surface of the capsid exhibiting high sequence diversity and variability may be linear epitopes for neutralizing antibodies; however, the proximity of some of these regions (Fig 8A, 8D, and 8G) could also form conformational epitopes. Clearly, high-resolution structures of mAb (or Fab) in complex with PCV2 are needed to differentiate these possibilities.

Currently used vaccines have been produced using the capsid from a PCV2a genome and several studies have reported immunization failures as a consequence of PCV2d infection (7, 48, 49). In addition, there is always the possibility of low vaccine efficacy due to genomic shift and this factor must be accounted for in future vaccine development. G. Franzo *et al* (39) studied the vaccine-derived selection pressure caused by vaccination. They reported that high mutation rates at amino acid positions 59, 191, 206 for PCV2a and 131, 228 for PCV2b reduced the binding of antibody that previously bound to the capsid; possibly causing the immune escape from vaccine protection (Fig 7) (50). Consequently, a platform for expression of PCV2d VLP is warranted.

The VLP expression system has multiple application. For example, the system can be used to generate capsids assembled as a multivalent mosaic that simultaneously displays neutralizing epitopes of several genotypes (i.e. transfect cells with multiple plasmids expressing different CP genotypes). Additionally, the 40 amino acids at the N-terminus can be replaced with specific tags to package biologics, expressed in the same cells, into the capsid. Such nanostructures could be utilized for diagnostics or delivery systems. There have been multiple reports demonstrating that VLP can assemble if the N-terminus of the CP is replaced with other sequences (11, 51).

In summary, we describe a mammalian assembly system of PCV2d VLPs, perform structural analysis based on cryo-EM image reconstruction, demonstrate a structural difference between the PCV2a, b, and d genotypes, model a tetranucleotide and a section of the N-terminus into the image reconstruction, demonstrate that the capsid packages nucleic acid, visualize the diversity and variability of the deposited PCV2 CP sequences on the CP structure, and identify conserved patches on the surface that may be taken advantage of for developing universal PCV2 vaccines.

## Materials and methods

### Cells, Capsid Gene, Plasmid and antibody

Suspension cultures of Expi293F human cells (Life Technologies, CA) were grown in serum-free Expi293 expression medium (Life Technologies, CA) at 37°C in a 5% CO_2_ environment and agitated at 150 rpm in Erlenmeyer flasks. The porcine circovirus type 2 (PCV2) gene encoding the capsid protein (GenBank: AWD32058.1) was chemically synthesized using a codon-optimized sequence by Blue Heron Technologies (Bothell, WA). The recognition site for NheI and the Kozak sequence were added right upstream from the start codon, and the recognition site for NotI was incorporated after the termination codon. The synthesized Cap gene was recovered from the transport plasmid by a double digestion with NheI and NotI restriction enzymes and sub-cloned after gel purification into the mammalian expression plasmid pcDNA3.4 cut with the same enzymes. The ligated plasmid was transformed into MAX Efficiency Stbl2 Cells (Life Technologies) and a correct clone was identified via restriction enzyme analysis and verified by sequencing.

### Virus Like Particle (VLP) Production and Purification

PCV2 VLPs were produced in a suspension culture of Expi293F mammalian cells following transient transfection with the plasmid pcDNA3.4-PCV2 (Fig. 1). Expi293F cells were seeded at the concentration of 2×10^6^ cells/ml and cultured for 16h prior to transfection. Plasmid DNA (1 µg/ml) was diluted in a volume of Opti-MEM representing 5% of the total volume of the culture. Separately, polyethylenimine (PEI) was prepared in an equivalent volume of Opti-MEM (4 µg/ml). After 5 min of incubation at room temperature, the PEI solution was added dropwise to the tube containing the DNA and after 30 min of incubation at RT the mixture was added to the cell suspension in a dropwise manner. Twenty-four hours after transfection, Valproic acid sodium salt (VPA) was added to the cell culture to a final concentration of 3.75 mM to inhibit cell proliferation. Seventy-two hours post-transfection the cells were pelleted by centrifugation at 2,000g for 15 min, and then washed one time with phosphate buffered saline (PBS) and spun again at 2,000g for 15 min. The cell pellet was re-suspended in PBS and then subjected to three freeze (−80°C) and thaw (37°C) cycles. Subsequently, the cells were further fragmented by three cycles of sonication and clarified by two successive centrifugations, first at 2,000g for 15 min followed by 8,000g for 15 min. The PCV2 VLPs contained in the clarified supernatant were further purified by ultracentrifugation on a two-layer CsCl density gradient: lower layer, 5 ml of 1.4 g/ml CsCl and upper layer, 10ml of 1.25 g/ml of CsCl both prepared in 10mM Tris-HCl, (pH 7.9). Samples were loaded onto the gradient and spun at 15°C for 4h at 140,000g using a SW28 rotor (Beckman Coulter, CA). The VLPs appeared as an opaque band at the interface of the 1.25 and 1.4 g/ml CsCl layers and were collected by piercing the tube with an 18G needle and syringe. The collected solution was mixed with 37 % CsCl in 10 mM Tris-HCl (pH 7.9) to final volume of 12 ml and then spun at 15°C for 16hr at 155,000g using a SW 41Ti rotor (Beckman Coulter, CA). The VLPs were detected at the lower part of the tube and recovered as described above. Collected VLP material was dialyzed against 10mM Tris-HCl pH 7.9 and 150mM NaCl at 10°C overnight using a Slide-A-Lyzer Cassette. Purified PCV2 VLPs were concentrated and buffer exchanged to phosphate buffered saline (PBS) using Amicon Utra-4 centrifugal filter devices (Merck Millipore, MA). PCV2 VLP samples were stored in 50-100 µl aliquots at −80°C.

### Western Blot and Coomassie Blue Stain

Purified VLPs were mixed with loading buffer, heated at 100°C for 5 min and run on a 4 to 12% Bis-Tris SDS-polyacrylamide gel (Life Technologies, CA). Loading amounts of proteins were 1 μg for Coomassie staining and 0.5 μg – for Western Blot. After electrophoretic separation, the gel was stained with Coomassie blue or proteins electro-transferred onto a 0.45 µm nitrocellulose membrane (Life Technologies LC2001). The membrane was then blocked with 5 % non-fat milk in TBST (10 mM Tris-HCl, 130 mM NaCl, and 0.05% Tween-20, pH 7.4) for 1h at (20°C) followed by an overnight incubation at (20°C) in primary Rabbit anti-porcine circovirus antibody (Cab 183908, Abcam, UK) diluted with blocking buffer. Membranes were washed 3 times with TBST and then incubated for 2 h with secondary antibody (goat anti-rabbit IgG HRP conjugated, 1:1,000) diluted in blocking buffer. Finally, membranes were washed 3 times with TBST and developed with ECL Western blot system (Life Technologies, CA) according to manufacturer’s instructions. The stained gel and immune blot images were acquired with a FluorChem Imager instrument (Protein Simple, CA).

### Immunofluorescence

The HEK293 adherent culture cells were transfected with pcDNA3.4-PCV2d plasmid using Amaxa Nucleofector II instrument by Lonza (program A-23). The transfected cell were plated on 6-well plate, each well contained cover slip to performed immunofluorescence study on transfected cells. The cover slips were removed 24, 48 and 72 hours post transfection and fixed with ice-cold acetone for 10 min. Expressed PCV2d CP was detected by using Mouse anti-PCV2 CP monoclonal antibody (GeneTex, GTX634211). The Donkey anti Mouse IgG conjugated with Alexa488 dye (Jackson ImmunoResearch) used as a secondary antibody. DAPI staining allowed us to localize the nuclei.

Images were acquired with a Neo sCMOS camera (6.45µm pixels, 560MHz, Andor Technology) on a Nikon TiE inverted microscope (Nikon Inc., Mellville, NY) using 40X (NA 0.95) plan apochromat objectives. 14-16bit images were scaled linearly to highlight features of interest and converted to 8-bit copies for figure assembly. Devices were controlled by Elements software (Nikon Instruments).

### Negative Staining and TEM Examination

5 uL of pCV2-2 VLP samples was applied to CF200-CU carbon film 200 mesh copper grids (Electron Microscopy Science) for 1 min and the grids were then washed with 200 µl of 50 mM of Na Cacodylate buffer and then strained immediately with 50 µl of 0.5% uranyl acetate for 1 min. The grids were examined by a JOEL 2100 transmission electron microscope operating at 200 kV with an Orius 2048 × 2048 pixel CCD (Gatan Inc., Pleasanton, CA).

### Cryo-EM Data Collection

Frozen hydrated samples of PCV2d VLPs were prepared on Quantifoil R 2/2, 200 mesh copper grids (Electron Microscopy Science). A 4 μl sample of the VLP was applied to the grid blotted for 3 seconds and flash frozen in liquid ethane using a FEI Vitrobot instrument. The grids were stored in liquid nitrogen until data collection. Data was collected at cryogenic temperatures on a FEI Titan Krios, operating at 300 kV, with Gatan K2 camera post a GIF quantum energy filter with a width of 15 eV. Data collection was performed with the Leginon suite (52).

### Image Reconstruction

The MotionCor2 package was used for correct for particle motion (53). Default parameters and dose weighting were used for the correction, with the patch 5 option, and the first frame of each movie was discarded during the alignment. The particles were selected automatically using Gautomatch v0.53, and contrast transfer function (CTF) estimation was performed on the aligned micrographs using Gctf v0.50 (54). Relion 3.0 was used to extract 23,358 300×300 pixel particles from the dose-weighted micrographs using the coordinates identified by Gautomatch. Reference free 2D classification was performed with Relion 3.0 (55). Non-default options for this step included a diameter of 250 Å and 128 classes were requested (56). An initial model was generated to 60 Å resolution using the PCV2 crystal structure (PDB entry 3R0R) with the *molmap* function of UCSF Chimera (57). 3D classification was carried out on 7196 particles using Relion 3.0 with a diameter of 250 Å, 3 classes, and C1 symmetry. A single class with 4,442 particles exhibited the highest resolution. These particles were used for a high-resolution image reconstruction with Relion 3.0. Again, a diameter of 250 Å and I1 symmetry were used with the remaining default parameters of Relion 3.0. A binary mask was created using the *relion_mask_create* program of Relion 3.0. The binary mask for postprocessing was generated as follows: 1) the high resolution image reconstruction was low pass filtered to 15 Å resolution using *relion_image_handler* (Relion 3.0), 2) the lowest threshold at which noise exterior to the PCV2 capsid was identified for this volume using UCSF Chimera, 3) *relion_mask_create* (Relion 3.0) was used to convert this volume into a binary mask with the identified threshold, mask dialation by 7 pixels and 2 soft edge pixels. The resulting mask was then inspected with UCSF Chimera to ensure that no internal cavities existed.

Local resolution was calculated with the program MonoRes. The same binary mask used during the postprocessing with Relion was used for the calculation. A resolution range of 3.3 Å to 5.3 Å was used (58).

### Structure refinement

The atomic coordinates for the crystal structure of PCV2b crystal structure (PDB entry 3R0R) were modified using Coot (59). The biological matrices necessary to generate a VLP are present in the PDB and were used by Coot to generate a VLP of PCV2d. UCSF Chimera was used to manually dock the VLP into the symmetrized image reconstructions. The resulting coordinates were iteratively refined using *phenix.real_space_refine* from the Phenix software package with non-crystallographic symmetry (NCS) constraints applied, and manual fitting with Coot (60).

### Sequence alignment, entropy calculation and evolution coupling

The nucleotide sequences used by Franzo and Segales were downloaded from GenBank (14). The NCBI ORFfinder suite was used to identify ORFs in each sequence. The ORFs identified from each entry was filtered to identify the CP ORF. The resulting ORFs were submitted to CD-HIT to identify a unique set of sequence for each genotype identified by Franzo and Segales (29). The sequence diversity and variability (entropy) for each genotype was calculated using the AL2CO routine built into the UCSF Chimera suite (57). The atomic coordinates of PCV2a, b, and d were used for the three genotypes studied (PDB entries 3JCI, 6DZU and 6OLA).

The PCV2 CP sequences were expanded by performing a protein Blast search with the sequence of PCV2d and the organism common name Porcine circovirus 2 (taxid:85708) filter (61). A total of 1,966 PCV2 sequences were identified. Partial sequences, sequences with names containing “putative”, “P3”, “unknown”, and “P27.9” were manually removed. Sequence alignment was performed on the remaining sequences using MUSCLE with default parameters (62). Sequences that generated gaps, possessed more than ten amino acids with a distinct sequence, or had no similar sequences were manually removed. This was done in order to remove spurious errors/artifacts that may have occurred during the sequencing process (i.e. artificial recombination during PCR with *Taq* polymerase (63)). Several rounds of alignment and deletion were performed to remove such sequences. The final set of sequences was combined with the pool of sequences identified by Franzo and Segales, and submitted to the CD-HIT server to attain a unique set of 1,278 sequences. The final round of alignment was performed with Clustal Omega with default parameters (64). Evolutionary coupling calculations could not be successfully performed with the EVcouplings server (evfold.org) because of the limited range in the Expect (E) value present the sequence (i.e. the sequences are too similar). Consequently, plmc was used to generate coevolution and covariation within the sequences. The L2 lambda for fields and couplings used were 0.01 and 16.0, respectively, and a maximum number of 100 iterations were performed. The results were converted for visualization with EVzoom using MATLAB 2019 and the scripts provided by the plmc program (33). The resulting matrix was visualized with EVzoom (33), and structural covariance was visually confirmed with UCSF Chimera (57).

### Measuring the interior content of the capsid

The radial profile of the image reconstruction was calculated using the *bradial* program from the Bsoft package (65). The radial profile in the interior of the capsid is quite strong and likely to arise from the PCV2 N-terminus (amino acids 1-36) and additional material. We assumed that cellular RNA was present in the capsid to stabilize the positive charge of the capsid interior (66). To calculate the amount of RNA present in the PCV2 capsid, we used the asymmetric image reconstruction of MS2 (EMD-8397) as a standard -as the ssRNA genome of MS2 is resolved in this image reconstruction (22). The intent is to adjust the threshold of the MS2 image reconstruction to account for 100% mass content in the capsid shell, then identify the threshold at which 100% mass content of the genome is accounted for. This threshold value is then used to measure the RNA content of the PCV2 image reconstruction after the PCV2 image reconstruction has been adjusted such that 100% mass content in the capsid shell is accounted for.

First, a 15 Å molecular envelope was generated for the MS2 capsid shell (PDB entry: 5TC1) with the UCSF Chimera *vop* command (57). The envelope was converted to the binary mask with the Relion 3.0 *relion_mask_create* program (55). The mask was applied to the image reconstruction using the Bsoft *bmask* program to extract the voxels pertaining to the capsid shell (65). The EMAN *volume* program was used to rescale the map densities to a threshold of 1.0 for a mass of 2.54 MDa - the total mass of 180 capsid proteins (*T=3*) and one maturation protein (22). The map densities of the unmasked image reconstruction was adjusted, via visual inspection with UCSF Chimera and scaling with Bsoft *bimg* program, such that its shell region was identical to the masked image reconstruction at threshold of 1.0 (2.54 MDa) (57, 65). The molecular envelope of the genome was isolated by applying an inverted mask of the capsid shell to remove the capsid shell, and a spherical mask to remove the solvent. The masks were applied with the Bsoft *bmask* program (65). The mass content of the remaining volume was calculated using the EMAN *volume* program (67). The Oligo Calc: Oligonucleotide Properties Calculator was used to determine the molecular weight of the MS2 genome (GenBank entry: NC_001417.2) to be 1.10 MDa (68). From this value we determined that at a threshold of 0.781 the MS2 genome is fully accounted for in the EMD-8397 image reconstruction whose map distribution has been adjusted to account for 100% mass content of the capsid shell.

Following a similar procedure for the PCV2 image reconstruction (amino acids 43-231), we determined that a threshold of 0.781 corresponds to a mass of 730 kDa in the capsid interior. From this mass we subtracted the mass of the 60 N-termini (amino acids 1-42, 5.8 kDa) that are believed to be inside the capsid and contribute to the envelope. If we assume that the remaining mass of 384 kDa is from RNA, this equates to approximately 1,200 nt. Approximate MW of ssRNA = (#nucleotides × 320.5) + 159.

### Measurement of neucleic acid content in purified VLP

The ratio of nucleic acid to protein (260nm:280nm) was determined after correction for scattering at 340nm and 360nm according to the protocol established by Porterfield and Zlotnick (23). Briefly, an adsorption scan of three sample were collected two times each from 640nm to 220nm using a VWR UV-1600 PC Spectrophotometer. The mass of the CP was calculated to be 28,057.04 Da using the amino acid sequence and the Expasy Protparam server (69). The extinction coefficient of the CP was calculated using the same server (70), which yielded a value of 48,360 M^−1^ cm^−1^. A 260 nm / 280 nm ratio for pure protein (0.6) was used to calculate the extinction coefficient for RNA 29,016 M^−1^ cm^−1^.

## Acknowledgement

Funds responsible for supporting these studies were provided by NIH National Institute of General medical Sciences and National institute of Allergy and Infectious Diseases (5SC1AI114843) to RK, by Grant Number 5G12MD007603-30 from the National Institute on Minority Health and Health Disparities to RK and PG, and by TechnoVax, Inc. and Huvepharma to JMG, KW, and BG. Data collection for the PCV2d VLP was performed at the Simons Electron Microscopy Center and National Resource for Automated Molecular Microscopy located at the New York Structural Biology Center, supported by grants from the Simons Foundation (349247), NYSTAR, and the NIH National Institute of General Medical Sciences (GM103310).

